# Sex differences in the amplification of responding to an alcohol-predictive cue by an alcohol-associated context

**DOI:** 10.1101/2020.09.10.292201

**Authors:** Diana Segal, Milan Valyear, Nadia Chaudhri

**Author notes:** Corresponding Author: Nadia Chaudhri, PhD, CSBN, Department of Psychology, Concordia University, 7141 Sherbrooke Street West, Room SP 244, Montreal, QC, H4B-1R6, Canada.

## Abstract

**Background:** In male rats, physical contexts that are associated with alcohol can invigorate responding to a discrete, alcohol-predictive conditioned stimulus (CS), and amplify priming-induced reinstatement. Here, we examined these effects as a function of biological sex.

**Methods:** Male and female Long-Evans rats were acclimated to drinking ethanol (EtOH, 15% v/v) in their home cages. Next, they were trained to associate an auditory CS (10 s; white noise; 15 trials per session) with EtOH delivery (0.2 ml per CS; 3.0 ml per session) into a fluid port for oral intake. Training occurred in a distinctive context containing specific visual, olfactory, and tactile stimuli. During alternating sessions rats were exposed to a second context where they did not receive EtOH. At test, CS presentations occurred in both contexts without EtOH delivery. Rats then underwent extinction using repeated unreinforced presentations of the CS in both contexts. An alcohol-primed reinstatement test was then conducted, in which 0.2 ml of EtOH was presented both at the start of the session and during the first CS presentation, after which no EtOH was delivered for the remainder of the session.

**Results:** At both test and reinstatement, male rats made significantly more CS port-entries in the context associated with alcohol delivery than in the context in which alcohol was never experienced. Unlike males, female rats made a similar number of CS port-entries at test in both the alcohol context and the neutral context. The reinstatement observed in female rats was not affected by context.

**Conclusions:** These findings identify novel sex differences in the capacity of an alcohol-associated context to modulate responding to a discrete, alcohol-predictive cue.

---

Almost half of the global population aged 15 years and older drinks alcohol, with increasing rates of problematic alcohol use producing serious consequences worldwide and contributing to about 3 million deaths every year (WHO, 2018). Alcohol use, misuse, and relapse in humans can be influenced by the environment in which alcohol is consumed (Ludwig et al., 1974; Ludwig,1986). Environmental stimuli can become associated with the pharmacological effects of alcohol, and trigger physiological, psychological and behavioural responses (Ludwig, 1986; Field and Duka, 2002; Powell, 2006). For example, exposure to a discrete alcohol-predictive conditioned stimulus (CS) (e.g. sound, sight, smell or taste) can induce craving in humans (Litt and Cooney, 1999; Witteman et al., 2015) and can reinstate extinguished alcohol seeking behaviour in animals (Bienkowski et al., 2000; Katner, 1999; Pina and Williams, 2016). Furthermore, environmental contexts that are associated with alcohol can facilitate craving in humans (Heinz et al., 2010; Ludwig et al., 1974; McCusker and Brown, 1990) and alcohol seeking in animals (Chaudhri et al., 2009, 2008a, 2008b; Janak, 2013; Marchant et al., 2013; Powell, 2006; Willcocks and McNally, 2011; Zironi et al., 2006). Discrete alcohol-predictive cues are often embedded within alcohol-associated contexts and the interaction between these types of stimuli may play a role in alcohol use and misuse.

We have previously assessed the interaction of an alcohol-associated context and a discrete alcohol-predictive CS, using a Pavlovian context discrimination task. Male rats were trained to discriminate between a context in which a discrete auditory CS was paired with alcohol delivery and a second different context in which a different auditory stimulus was presented without alcohol delivery. We found that responding to a discrete alcohol-predictive CS was amplified in the alcohol-associated context, compared to the context in which alcohol was never consumed (Khoo et al., 2019a, 2019b; Millan et al., 2015; Remedios et al., 2014; Sciascia et al., 2015; Valyear et al., 2020). This result indicates that conditioned responding to an alcohol-predictive CS is context-dependent in male rats, but the context-dependency of conditioned responding to an alcohol-predictive CS has not been assessed in females.

Studies assessing sex differences in context-dependent learning suggest that females may not show context-induced amplification of conditioned responding. Evidence from preclinical studies suggests that males and females differ in their dependency on the environment and/or discrete cues. For example, sex differences in context-modulation of cue-elicited responding exist in appetitive ABA renewal. At test males in the experimental group (i.e. ABA) display more cue-elicited responding compared to male controls (i.e. AAA), while females in both groups respond similarly during the cue (Anderson and Petrovich, 2015, 2017, 2018a, 2018b). Females are also more likely to generalize conditioned responding to novel contexts. In contextual fear conditioning, females display higher levels of freezing in novel contexts, compared to males. However, following context preexposure, context discrimination increases in females (Asok et al., 2019; Keiser et al., 2017; Lynch et al., 2013; Wiltgen et al., 2001). Thus, males may be more efficient in forming contextual representations than females. Furthermore, studies using spatial tasks suggest that males and females use different contextual information to navigate through their environment. In tasks like radial arm mazes performance in males is disrupted by changes to the euclidean (i.e. distance and angle) and geometric properties of the environment, while in females performance is disrupted by changes in landmark cues (i.e. objects in a room) (Rodríguez et al., 2011; Tropp and Markus, 2001; Williams et al., 1990; Williams and Meck, 1991). Therefore, sex differences in the utilization of contextual information may impact how alcohol-associated contexts modulate responding elicited by alcohol-predictive cues in males and females.

Similarly, studies in humans have suggested that men and women use contextual information differently. Men have been shown to predominately use their environment to complete navigation tasks or virtual mazes and typically complete the task faster than women, who predominantly use landmark cues. However, these sex differences in performance are not evident when tasks can be completed using landmark information (Boone et al., 2018), which suggests that men and women learn about their environments differently and use different types of contextual cues to complete tasks. Thus, the differences in the way men and women learn and associate contextual information may influence how context modulates their behaviour in situations involving substance use. However, studies investigating how environmental contexts modulate alcohol use in men and women are limited. Recently, it was shown that both men and women with alcohol use disorder report high levels of alcohol craving in alcohol-associated contexts (e.g. a bar, a pub and parties), but women also reported higher levels of cravings in contexts not commonly associated with alcohol (e.g. the workplace, bedroom and supermarket) (Ghiţă et al., 2019). Thus, alcohol craving in men may be more specific to certain environmental contexts than in women. However, it is not known whether alcohol-associated contexts influence responding to discrete alcohol-predictive cues similarly in women and men.

In addition to sex differences in context-dependent learning, both human and animal studies suggest that females may be more vulnerable to drug misuse and relapse (Hudson and Stamp, 2011; Zlebnik, 2019). Female rats acquire self-administration of drugs at a faster rate, escalate drug taking more rapidly, and show greater reinstatement of drug-seeking, compared to males (Becker and Koob, 2016). The importance of assessing gender differences in the mechanisms promoting alcohol misuse is further highlighted by recent evidence in clinical studies. In recent decades, problematic alcohol use in women has drastically increased, from adolescence through older adult years (Becker et al., 2017; Keyes et al., 2019). Women are more sensitive to the pharmacological effects of alcohol (Miller et al., 2009) and are at a greater risk for alcoholism-related cognitive and bodily impairments, than men (Becker et al., 2017; Ceylan-Isik et al., 2010). Therefore, there is an increasing need for research directed towards understanding the influence of context on CS-induced alcohol use in both men and women.

In the present study we examined the influence of an alcohol-associated context on responding to a discrete, alcohol-predictive CS in male and female rats. Rats were trained to associate a discrete auditory CS with alcohol delivery into a fluid port in a multimodal context. On alternating days, rats were placed into a different neutral context in which a different auditory stimulus was presented without the delivery of alcohol. Entries into the fluid port during the CS were assessed as the main dependent variable. Following Pavlovian discrimination training, CS port-entries were tested in both contexts without alcohol delivery, in order to examine the influence of context on responding to a CS. Next, CS-elicited port-entries were extinguished in both contexts, and then we examined the impact of context in an alcohol-primed induced reinstatement test. Based on prior results (Khoo et al., 2019a, 2019b; Millan et al., 2015; Remedios et al., 2014; Sciascia et al., 2015; Valyear et al., 2020), we predicted that males would show invigoration of CS-elicited port-entries in the alcohol-associated context compared to the neutral context, during the test and alcohol-primed reinstatement test. Based on growing evidence described above that behaviour in females is not context-dependent, we predicted that responding would not differ between the alcohol-associated context and the neutral context in female rats.

## MATERIALS AND METHODS

### Animals

Thirty-one Long-Evans rats (n = 16 males, n = 15 females; bred in-house; see Supplementary Material) were used. Rats were single housed in polycarbonate shoebox cages containing beta chip bedding (Aspen Sani chips; Envigo, Indianapolis, IN), a nylabone^™^ chew toy (Nylabones, Bio-Serv, Flemington, NJ), a red plastic tunnel (Rat retreats, Bio-Serv, Flemington, NJ), and shredded paper for enrichment, with unrestricted access to standard rat chow (Rodent Diet, Charles River, St. Hubert, QC) and water. Male and female cages were housed side by side in a colony room held at 21 +/- 2°C and approximately 40-50% humidity on a 12-hour light-dark cycle (lights on at 0700 hours). All procedures were conducted in the light phase. Rats were given 1 week to acclimate and 1 week to be handled by the experimenter before starting behavioural testing. All experimental procedures were approved by the Animal Research Ethics Committee at Concordia University and complied with regulations provided by the Canadian Council on Animal Care.

### Apparatus

Twenty-two conditioning chambers (ENV-009A) enclosed in fan-ventilated (72-80 dB background noise) sound-attenuating melamine cubicles (53.6 ⨯ 68.2 ⨯ 62.8 cm) from Med-Associates Inc (St. Albans, VT, USA) were used for behavioural training and testing. Conditioning chambers were made of stainless-steel bar floors (ENV-009A-GF), paneled aluminum sidewalls, and clear polycarbonate rear walls, ceilings, and front doors. Each chamber featured a white house-light (ENV-215M), and a white noise (∼8 dB above background; ENV-225SM) and clicker (5 Hz, ∼8 dB above background; ENV-135M) generator on the upper left wall. The right wall of each chamber contained a dual fluid port (ENV-200R3AM). A syringe pump (PHM-100, 3.33 rpm) located outside the cubicle that was used to deliver alcohol via a 20 ml syringe into the fluid well within the chamber. The entrance of the fluid port contained an infrared photobeam (ENV-205M) that when transected was counted and recorded as a port-entry on a PC computer. House light illumination, stimulus presentations, and fluid delivery were controlled by Med PC-IV software.

### Solutions

Ethanol (EtOH; 5, 10, 15%; *v/v*) was prepared weekly by diluting 95 % EtOH in tap water. Lemon oil (used as lemon odor; SAFC Supply Solutions, St-Louis, USA) or benzaldehyde (used as almond odor; ACP Chemicals Inc., Montreal, Canada) were mixed in tap water to obtain 10% solutions (*v/v*).

### Home-Cage Ethanol Exposure

To acclimate rats to drinking ethanol, a total 15 EtOH sessions were conducted. Every other day (i.e. Monday, Wednesday, Friday) rats were given 24-hour access to 15% EtOH and water via a 100 ml graduated cylinder and 500 ml bottle that were placed onto the home cage lids (Maddux, Lacroix, & Chaudhri, 2014; Simms et al., 2008; Wise, 1973). On Tuesday, Thursday, Saturday, and Sunday, EtOH cylinders were replaced with water bottles (procedure adapted from Sparks, Sciascia, Ayorech, & Chaudhri, 2014). The position (left or right) of the EtOH cylinders and water bottles were alternated to reduce the impact of side preference. At the end of each session containers were weighed. To control for spillage and evaporation, two empty control cages were set up on the highest and lowest shelves containing rats and were treated identically to the home cages. The average volume of fluid lost from the bottles in the cages were subtracted from the bottles in the home cages for each corresponding session. The grams of ethanol and water and the ingested dose of ethanol (g/kg; grams of ethanol consumed accounting for the density, per kg of body weight) were recorded for each rat in each 24-hour session.

Rats that consumed <1 g/kg of 15% EtOH averaged across sessions 5 and 6, were given access to 5% EtOH to encourage intake (Cofresí et al., 2017). When they reached ≥ 1 g/kg averaged across two consecutive sessions, they were given access to 10% EtOH until they once again reached ≥ 1 g/kg averaged across two consecutive sessions, after which they were given access to 15% EtOH. Rats that did not consume an average of ≥ 1 g/kg remained on the last given percentage of EtOH for the remainder of the experiment. In total, twenty-three rats (male = 13, female = 10) were trained and tested using 15% ETOH, the remaining eight were trained and tested using either 10% EtOH (males = 2, females = 1) or 5% EtOH (male = 1, female = 4).

### Pavlovian Discrimination Training

#### Habituation

During the last week of the home-cage ethanol exposure, 3 habituation sessions were conducted on days that rats only had access to water. During the first session, rats were brought to the experimental room in their home cages, weighed, and left for 20 mins, in order to habituate them to the new environment. On the following 2 sessions, rats were habituated to the conditioning chambers in both contexts, Context A on the first day and Context B on the second day in a counterbalanced design (See Table 1 for description of contexts). During these sessions, rats were weighed and placed in the conditioning chambers, and after a 1 min delay the house-light turned on for 20 minutes and entries into the fluid port were recorded in each session.

**Table 1.** Description of contexts used for Pavlovian discrimination training.

| Modality | Context A | Context B |
| --- | --- | --- |
| Visual | Black cardboard covered walls | Clear plexiglass walls |
|  | Brown paper in waste tray | Brown paper in waste tray |
| Tactile | Clear polycarbonate floor | Wire-grid floor |
| Olfactory<sup>1</sup> | Lemon odour | Almond odour |

<sup>1</sup> Sprayed (3 sprays) onto petri dish placed in the waste tray beneath the chamber.

#### Training

Rats underwent Pavlovian conditioning with context alternation of an alcohol context and a neutral context (Figure 1A) that occurred continuously from the first to the last session of the experiment. Rats were counterbalanced across contexts and stimuli such that there were no differences in home-cage ethanol consumption. Conditioning chambers were designated based on sex, so that both sexes were never placed into the same chamber. Rats were given one training session a day (Monday to Friday) until they had received 12 sessions in each context.

**Figure 1.**
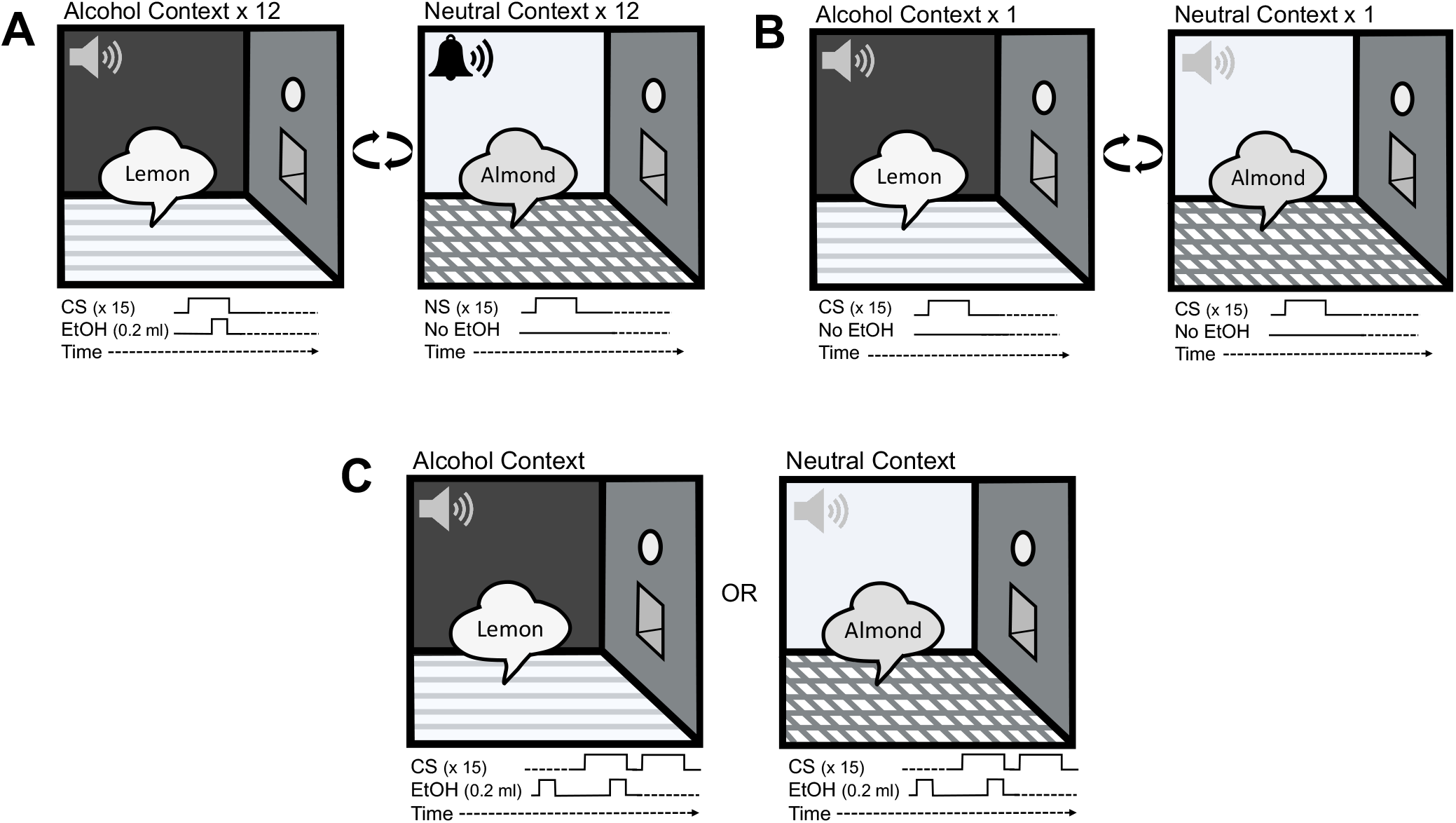
Schematic of Pavlovian discrimination paradigm. (**A**) During Pavlovian discrimination training, rats received 24 sessions (12 in each context), in which a discrete auditory conditioned stimulus (CS) was paired with 15% ethanol (EtOH). On alternating days, rats were exposed to a neutral context where a neutral auditory stimulus (NS) was presented without EtOH. (**B**) At test, the CS was presented in both contexts, but no alcohol was delivered. (**C**) Following extinction using repeated tests, a between-subjects alcohol-primed reinstatement test was conducted where the CS was presented in both contexts and 0.2 ml of EtOH was delivered at the start of the session and during the first CS presentation, after which no EtOH was delivered for the remainder of the session.

Training sessions (73.5 mins duration) began with a two-minute delay, after which the house-light turned on and the first inter-trial interval (ITI) began. Sessions conducted in the alcohol context consisted of 15 trials of an auditory CS (10 s; continuous white noise or clicker) that occurred on a variable-time 240 s schedule. Prior to and following each CS interval was a 10 s PreCS and PostCS interval. Therefore, between the PostCS offset and subsequent PreCS onset, an ITI occurred for either 120, 240, or 360 s. Every CS presentation was paired with 0.2 ml of 15% ethanol dispensed into the fluid port over the last 6 s of the CS (total of 3 ml of ethanol per session). At the end of each session, all the ports were checked to ensure that the ethanol was consumed. Sessions conducted in the neutral context were similar to those in the alcohol context with the exceptions that a different auditory neutral stimulus (NS; continuous white noise or clicker) was presented to equate acoustic valence of the context, and no alcohol was dispensed into the fluid ports. Syringe pumps were activated on a similar schedule, but they did not contain any syringes.

Rats that made < 15 CS port-entries averaged across sessions 5 and 6, in the alcohol context (male=4, female=2), were given a 2% sucrose and ethanol solution (15%=5, 5%=1) during subsequent training sessions. Once an average of ≥ 15 CS port-entries across two consecutive sessions was reached, rats were switched back to the ethanol solution without sucrose for the remainder of the experiment.

#### Test: Effect of Context on Responding to CS

Following the last training session, the effect of context on CS-elicited port-entries was tested, using a counterbalanced within-subjects design (Figure 1B). Two test sessions where conducted using the procedure as in training, with the exception that all the rats received their CS without ethanol delivery. The test sessions were separated by 4 re-training sessions (2 in each context).

### Alcohol-Priming Induced Reinstatement Test

Following the last test session, 4 re-training sessions were conducted (2 in each context). Next, a total of 8 repeated test sessions (4 in each context) were conducted to extinguish CS-elicited port-entries in the alcohol context and the neutral context.

After the last repeated test session, an alcohol-priming induced reinstatement test was conducted using a between-subjects design (Figure 1C). At test, 0.2 ml of ethanol was dispensed over 6 s into the fluid port, 30 s after the start of the session. Additionally, 0.2 ml of ethanol was dispensed over the first 6 s during the first CS trial, after which no ethanol was dispensed for the remainder of the session. The ethanol prime served as a reminder of the orosensory properties of the ethanol. The reinstatement test was conducted in the opposite context from that in the last repeated test session. Therefore, rats that completed the last repeated test session in their alcohol context, received the reinstatement test in their neutral context (male = 8, female = 8) and vice versa (male = 8, female = 7).

### Statistical Analysis

Five (2 = males, 3 = females) of the original 31 rats that underwent Pavlovian discrimination training were excluded from the statistical analysis as they did not meet acquisition criteria. These criteria were 10 or more CS port-entries and a > 50% probability of making at least 1 port-entry per trial, averaged across the last two alcohol sessions.

During the Pavlovian discrimination training, entries into the fluid port during the 10 s PreCS/NS, 10 s CS/NS, and total port-entries were measured in each session. Additionally, the total latency to initiate CS port-entries and the total duration of all CS port-entries were recorded. A ΔCS port-entry (CS minus PreCS) variable was calculated for the reinstatement test to account for individual differences in baseline port-entries. Responding at the reinstatement test was compared to the extinction baseline (average ΔCS across the last two extinction sessions).

The interquartile range method (Tukey, 1977) was used to identify and correct outliers by replacement with the median. Three data points were identified as outliers; the duration of CS port-entries from a female rat (session 7 of training; neutral context), the duration and number of CS port-entries from a female rat (session 4 of extinction; alcohol context).

Data were analyzed using mixed ANOVAs, with a Greenhouse-Geiser correction applied when Mauchly’s test of sphericity was significant. Our prior research has reliably found that male rats make more CS port-entries at test in the alcohol context than in the neutral context (Millan et al., 2015; Remedios et al., 2014; Sciascia et al., 2015; Valyear et al., 2020). Therefore, planned comparisons were conducted to analyze CS port-entries across contexts during test and the reinstatement test separately in both sexes, using paired- or independent-samples t-tests with Cohen’s d reported as the effect size. Analyses were conducted with JASP Statistics version 0.11.1 (JASP Team, University of Amsterdam, NL) and graphs were created with Prism 8 (GraphPad Statistics, La Jolla, CA).

## RESULTS

### Home-Cage Ethanol Exposure

Males weighed more than females throughout home-cage ethanol exposure (Figure 2A). The weight (g) of males (n=14) and females (n=12) increased across sessions [Session, *F*_(1.54, 37.0)_=263.45, *p*=< .001, *η*^*2*^=.038] and was greater in males compared to females [Sex, *F*_(1,24)_=146.92, *p*=< .001, *η*^*2*^=.860; Session x Sex, *F*_(1.54, 37.0)_=36.54, *p*=< .001, *η*^*2*^=.005].

**Figure 2.**
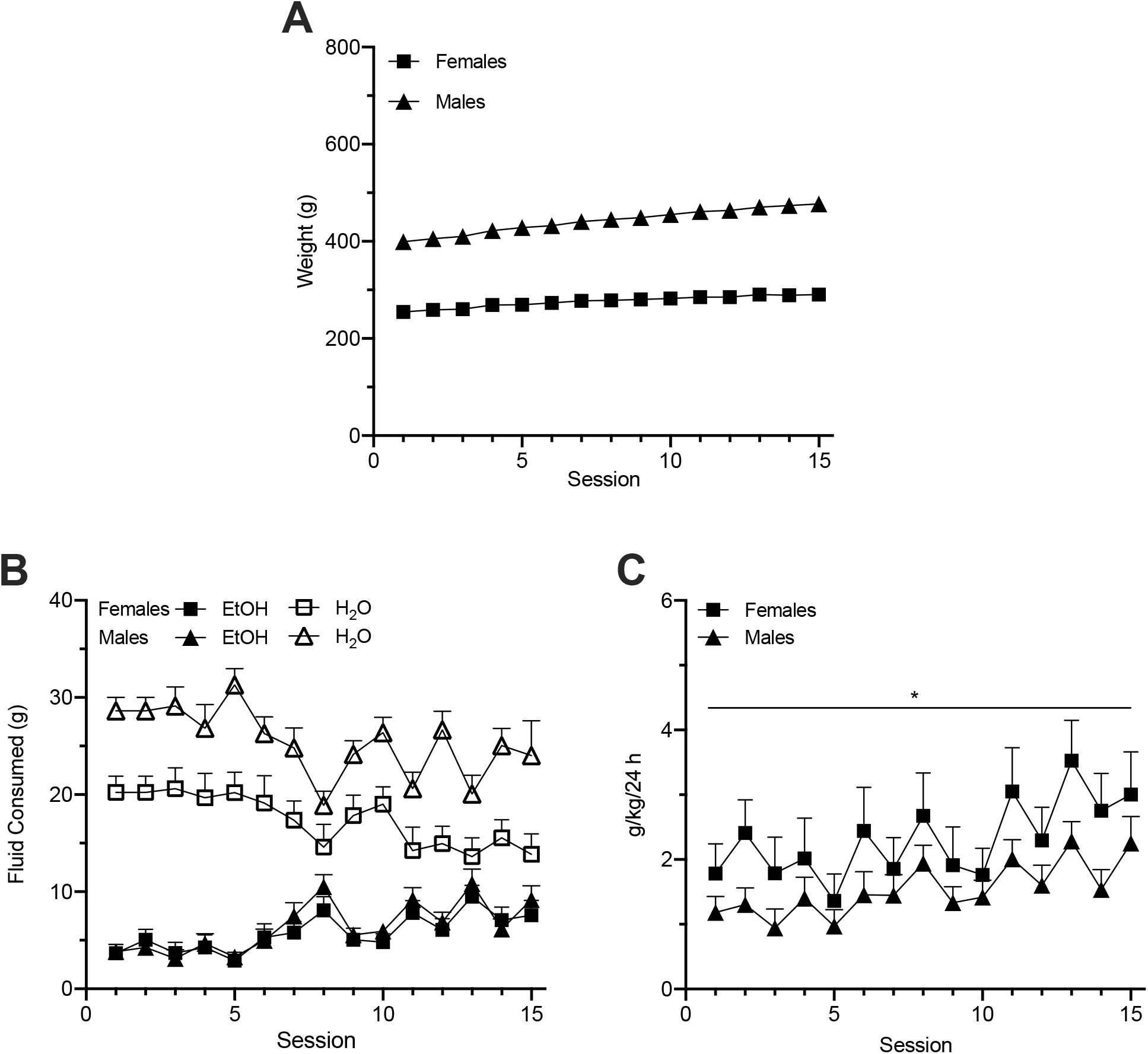
Intermittent access to alcohol and water in the home-cage in males (triangles) and females (squares). (**A**) Mean (± SEM) weight (g) of rats during the home-cage ethanol exposure. Males weighed more than females across sessions. (**B**) Mean (± SEM) grams of ethanol (filled symbols) consumed in each 24-h session was similar across sex, while mean (± SEM) grams of water (empty symbols) consumed in each 24-h session was greater in males compared to females. (**C**) Mean (± SEM) ingested dose of ethanol (grams of ethanol per kilogram of bodyweight accounting for the density of ethanol) per 24-h session was comparable in both males and females.

Analysis of fluid intake revealed that males drank more water (g) than females, whereas ethanol consumption (g) was similar in both sexes (Figure 2B). Overall, rats drank more water than ethanol [Fluid, *F*_(1,24)_=88.28, *p* < .001, *η*^*2*^=.451] and males consumed more fluid than females [Sex, *F*_(1,24)_=25.03, *p*<.001, *η*^*2*^ =.511]. The ANOVA revealed a significant Fluid x Sex interaction [*F*_(1, 24)_=6.08, *p*=.021, *η*^*2*^=.031]. Follow-up independent samples t-tests indicated no difference in ethanol intake in males (*M*=6.39 ± *SE*=.69) and females (*M*=5.77 ± *SE*=.864) [*t*_(24)_=0.56, *p*=.582, *d*=0.22, 95% CI (-.556, .991)], but greater water intake in males (*M*=25.86 ± *SE*=1.66) than females (*M*=17.15 ± *SE*=1.78) [*t*_(24)_=3.58, *p*=.002, *d*=1.41, 95% CI (.529, 2.263)].

Overall, fluid intake did not vary across sessions [Session, *F*_(1.33, 31.81)_=0.78, *p*=.419, *η*^*2*^=.003] in either sex [Session x Sex, *F*_(1.33, 31.81)_=0.93, *p*=.370, *η*^*2*^=.004]. Water and ethanol intake changed from the first to last sessions [Fluid x Session, *F*_(2.78, 66.82)_=7.12, *p* < .001, *η*^*2*^=.046] comparably in both sexes [Fluid x Session x Sex, *F*_(2.78, 66.82)_=0.58, *p*=.617, *η*^*2*^=.004]. Collapsed across sex, ethanol intake increased from session 1 (*M*=3.75 ± *SE*=0.58) to session 15 (*M*=8.44 ± *SE*=1.04) [*t*_(25)_=-4.89, *p* < .001, *d*=-0.96, 95% CI (−1.419, −0.486)], whereas water intake remained similar from session 1 (*M*=24.76 ± *SE*=1.34) to session 15 (*M*=22.80 ± *SE*=5.84) [*t*_(25)_=0.35, *p*=.731, *d*=0.07, 95% CI (−0.32, 0.45)].

Males and females did not differ in overall ingested dose of ethanol throughout the home-cage ethanol exposure phase (Figure 2C; [Sex, *F*_(1,24)_=2.117, *p*=.159, *η*^*2*^=.081]. The ingested dose of ethanol varied as a function of session [Session, *F*_(4.77, 114.47)_=9.00, *p* < .001, *η*^*2*^=.085] similarly in both sexes [Session x Sex, *F*_(4.77, 114.47)_=0.80, *p*=.550, *η*^*2*^=.008]. Follow-up paired samples t-tests on the data collapsed across sex indicated that the ingested dose in session 15 (*M*=2.60 ± *SE*=.38) was higher than in session 1 (*M*=1.46 ± *SE*=0.25) [*t*_(25)_=3.72, *p*=.001, *d*=0.73, 95% CI (0.29, 1.16)].

### Pavlovian Discrimination Training

Both males and females learned to associate a CS with ethanol in the alcohol context, with females making more CS port-entries than males (Figure 3A). ANOVA comparing port-entries made during the PreCS or PreNS and CS or NS intervals in the alcohol and neutral contexts indicated that overall, port-entries varied as a function of session [Session, *F*_(3.89, 93.27)_=8.45, *p* < .001, *η*^*2*^=.031], were higher in the alcohol context than in the neutral context [Context, *F*_(1,24)_=130.86, *p* < .001, *η*^*2*^=.145], were higher in females than males Sex, *F*_(1, 24)_=5.14, *p*=.033, *η*^*2*^=.176] and varied as a function of interval [Interval, *F*_(1,24)_=125.79, *p* < .001, *η*^*2*^=.174]. Across sessions, CS port-entries increased, whereas port-entries during the NS, PreCS, and PreNS intervals remained low [Session x Interval, *F*_(4.28, 102.61)_=14.98, *p*<.001, *η*^*2*^=.040; Context x Session, *F*_(4.13, 101.23)_=12.96, *p*<.001, *η*^*2*^=.037; Context x Interval, *F*_(1,24)_=140.85, *p*<.001, *η*^*2*^=.102; Context x Interval x Session, *F*_(4.64, 111.44)_=12.10, *p*<.001, *η*^*2*^=.029]. A significant Context x Interval x Sex interaction [*F*_(1,24)_=4.89, *p*=.037, *η*^*2*^=.004] was found. Follow-up Bonferroni-corrected independent sample t-tests indicated that collapsed across sessions, in the alcohol context females made more CS port-entries than males [*t*=-4.15, *p*=.002, 95% CI (−6.00, −10.67)]. There was no sex difference in PreCS port-entries in the alcohol context [*t*=-0.80, *p* > .999, 95 CI (−5.82, 3.52)] or in NS [*t*=-0.49, *p* >.999, 95% CI (−5.372,3.965)] or PreNS [*t*=-0.36, *p* >.999, 95% CI (−5.19, 4.14)] port-entries in the neutral context. No other sex differences were found [Session x Sex, *F*_(3.89, 93.27)_=0.77, *p*=.542, *η*^*2*^=.003; Interval x Sex, *F*_(1,24)_=2.98, *p*=.097, *η*^*2*^=.004; Context x Sex, *F*_(1,24)_=5.15, *p*=.033, *η*^*2*^=.006; Session x Interval x Sex, *F*_(4.28, 102.61)_=0.75, *p*=.571, *η*^*2*^=.002; Context x Session x Sex, *F*_(4.22, 101.22)_=0.84, *p*=.510, *η*^*2*^=.002; Context x Interval x Session x Sex, *F*_(4.64, 11.44)_=0.82, *p*=.532, *η*^*2*^=.002].

**Figure 3.**
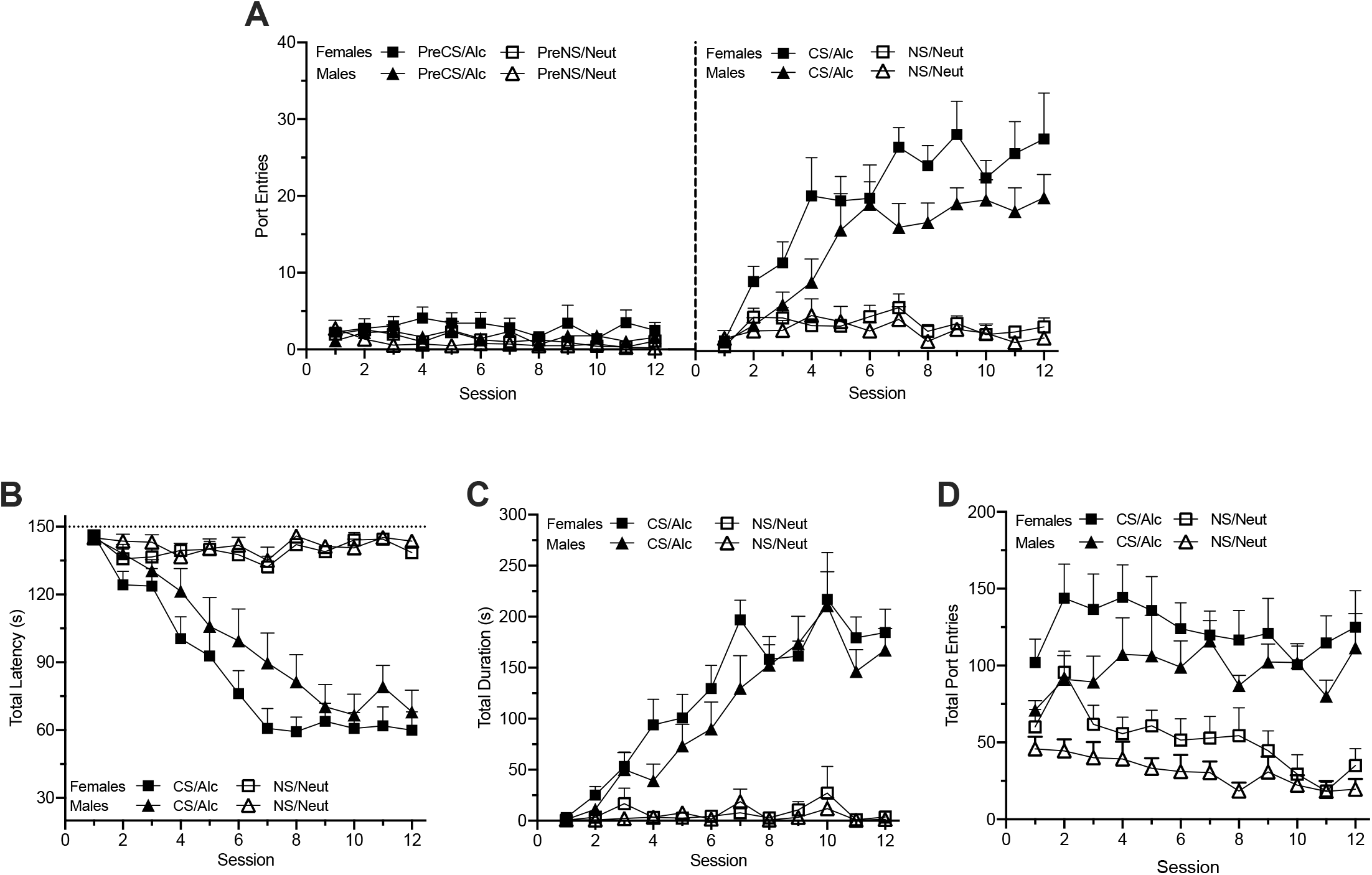
Acquisition of Pavlovian discrimination training in the alcohol context (filled shapes) and neutral context (empty shapes) in males (triangles) and females (squares). (**A**) Mean (± SEM) port entries during the PreCS or PreNS (left panel) and CS or NS (right panel) intervals. CS port entries were greater than NS, PreCS, and PreNS port entries in both sexes, with females making more CS port entries than males. (**B**) Mean (± SEM) total latency (s) to initiate port entries during the CS or NS interval. The total latency of CS port entries in the alcohol content was faster than the total latency of NS port entries in neutral context comparably in both sexes. (**C**) Mean (± SEM) total duration of port entries initiated during the CS or NS interval. The total duration of CS port entries in the alcohol context was greater than in the total duration of NS port entries in the neutral context comparably in males and females. (**D**) Mean (± SEM) total port entries (i.e. all the port entries in each session). Total port entries were greater in females than in males, remained high in the alcohol context, and decreased in the neutral context across sessions.

The total latency to initiate a port-entry after stimulus onset was shorter for the CS in the alcohol context compared to the NS in the neutral context in both males and females (Figure 3B). Overall the total latency to initiate a CS or NS port-entry varied as a function of session [Session, *F*_(4.45, 106.68)_=36.88, *p*<.001, *η*^*2*^=.127] in the alcohol context but not in the neutral context [Context, *F*_(1,24)_=109.77, *p*<.001, *η*^*2*^=.261; Context x Session, *F*_(5.35, 128.42)_=43.43, *p*<.001, *η*^*2*^=.124]. Follow-up paired samples t-tests collapsed across sex revealed that the latency to make a CS port-entry in the alcohol context decreased from session 1 (*M*=145.53 ± *SE*=1.25) to session 12 (*M*=64.35 ± *SE*=6.29) [*t*_(25)_=12.33, *p* < .001, *d*=2.42, 95% CI (1.64, 3.18)], whereas latency to make a port-entry following NS onset in the neutral context remained stable and high [*t*_(25)_=1.59, *p*=.124, *d*=.31, 95% CI (−0.09, 0.70)] from session 1 (*M* = 145.46 ± *SE* = 1.76) to session 12 (*M* = 141.28 ± *SE* = 2.04). No differences across sex were found [Sex, *F*_(1,24)_=1.75, *p*=.198, *η*^*2*^=.068; Session x Sex, *F*_(4.45, 106.68)_=1.05, *p*=.390, *η*^*2*^=.004; Context x Sex, *F*_(1,24)_=1.60, *p*=.217, *η*^*2*^=.005; Context x Session x Sex, *F*_(5.35, 128.42)_=1.06, *p*=.387, *η*^*2*^=.003].

The total duration of port-entries initiated during the CS was longer and increased over time in the alcohol context, while the total duration of port-entries initiated during the NS in the neutral context remained long, in both males and females (Figure 3C). Overall, total duration of port-entries initiated during the CS or NS varied across sessions [Session, *F*_(2.90, 69.55)_=26.06, *p*<.001, *η*^*2*^=.137], and was greater in the alcohol context than in the neutral context [Context, *F*_(1,24)_=111.50, *p*<.001, *η*^*2*^=.324] and varied as a function of session in the alcohol context but not in the neutral context [Context x Session, *F*_(2.88, 69.01)_=20.82, *p*<.001, *η*^*2*^=.107]. Follow-up paired samples t-tests collapsed across sex revealed that the total duration of CS port-entries in the alcohol context increased from session 1 (*M* = 2.05 ± *SE* = .85) to session 12 (*M* = 175.31 ± *SE* = 15.27) [*t*_(25)_=-11.31, *p*<.001, *d*=-2.22, 95% CI (−2.03, −1.49)], while the total duration of NS port-entries in the neutral context remained low [*t*_(25)_=-1.53, *p*=.139, d=-0.30, 95% CI (−0.69, 0.10)] from session 1 (*M* = 0.80 ± *SE* = .45) to 12 (*M* = 2.44 ± *SE* = .95). No differences across sex were found [Sex; *F*_(1,24)_=1.18, *p*=.289, *η*^*2*^=.005; Context x Sex, *F*_(1,24)_=0.70, *p*=.413, *η*^*2*^=.002; Context x Session x Sex, *F*_(2.88, 69.01)_=1.24, *p*=.301, *η*^*2*^=.006].

Total port-entries remained high in the alcohol context, and decreased in the neutral context throughout training, with females making more port-entries than males (Figure 3D). Overall, total port-entries were greater in females than in males [Sex, *F*_(1,24)_=4.35, *p*=.048, *η*^*2*^=.034], varied across sessions [Session, *F*_(4.95, 116.34)_=2.37, *p*=.046, *η*^*2*^=.023], were greater in the alcohol context than in the neutral context [Context, *F*_(1,24)_=104.06, *p*<.001, *η*^*2*^=.290], and varied as a function of session in the neutral context but not in the alcohol context [Context x Session, *F*_(6.12, 146.92)_=2.30, *p*=.036, *η*^*2*^=.013]. Follow-up paired samples t-tests collapsed across sex revealed that total port-entries in the alcohol context remained high [*t*_(25)_=-1.81, *p*=.083, d=-0.35, 95% CI (−0.75, 0.05)] from session 1 (*M* = 85.35 ± *SE* = 8.20) to session 12 (*M* = 117.73 ± *SE* = 16.00), while total port-entries in the neutral context decreased [*t*_(25)_=3.19, *p*=.004, d=0.63, 95% CI (0.20, 1.04)] from session 1 (*M* = 52.46 ± *SE* = 6.72) to session 12 (*M* = 26.69 ± *SE* = 6.27). No other sex differences were found [Context x Sex, *F*_(1,24)_=0.21, *p*=.653, *η*^*2*^<.001; Session x Sex, *F*_(4.85, 116.34)_=0.92, *p*=.466, *η*^*2*^=.009; Context x Session x Sex, *F*_(6.12, 146.92)_=0.57, *p*=.756, *η*^*2*^=.003].

### Test: Effect of Context on Responding to CS

Following Pavlovian discrimination training, CS port-entries were tested in the absence of alcohol delivery in both the alcohol context and in the neutral context. CS port-entries were higher in the alcohol context compared to the neutral context in males, while females responded similarly to the CS in both contexts (Figure 4A). ANOVA was conducted to assess PreCS and CS port-entries at test in both contexts and sexes. Overall, responding was greater during the CS than the PreCS interval [Interval, *F*_(1,24)_=69.74, *p*<.001, *η*^*2*^=.495]. There was no difference in the overall number of port-entries as a function of sex [Sex, *F*_(1,24)_=0.49, *p*=.491, *η*^*2*^=.020] or context [Context, *F*_(1,24)_=1.06, *p*=.313, *η*^*2*^=.003]. CS port-entries were higher than PreCS port-entries in both males and females [Interval x Sex, *F*_(1,24)_=0.52, *p*=.477, *η*^*2*^=.004] and there was a trending Context x Sex interaction [*F*_(1,24)_=3.83, *p*=.062, *η*^*2*^=.011]. There was no difference in the number of port-entries made in each context as a function of interval [Context x Interval, *F*_(1,24)_=0.55, *p*=.466, *η*^*2*^=.002; Context x Interval x Sex interaction, *F*_(1,24)_=3.14, *p*=.089, *η*^*2*^ =.009].

**Figure 4.**
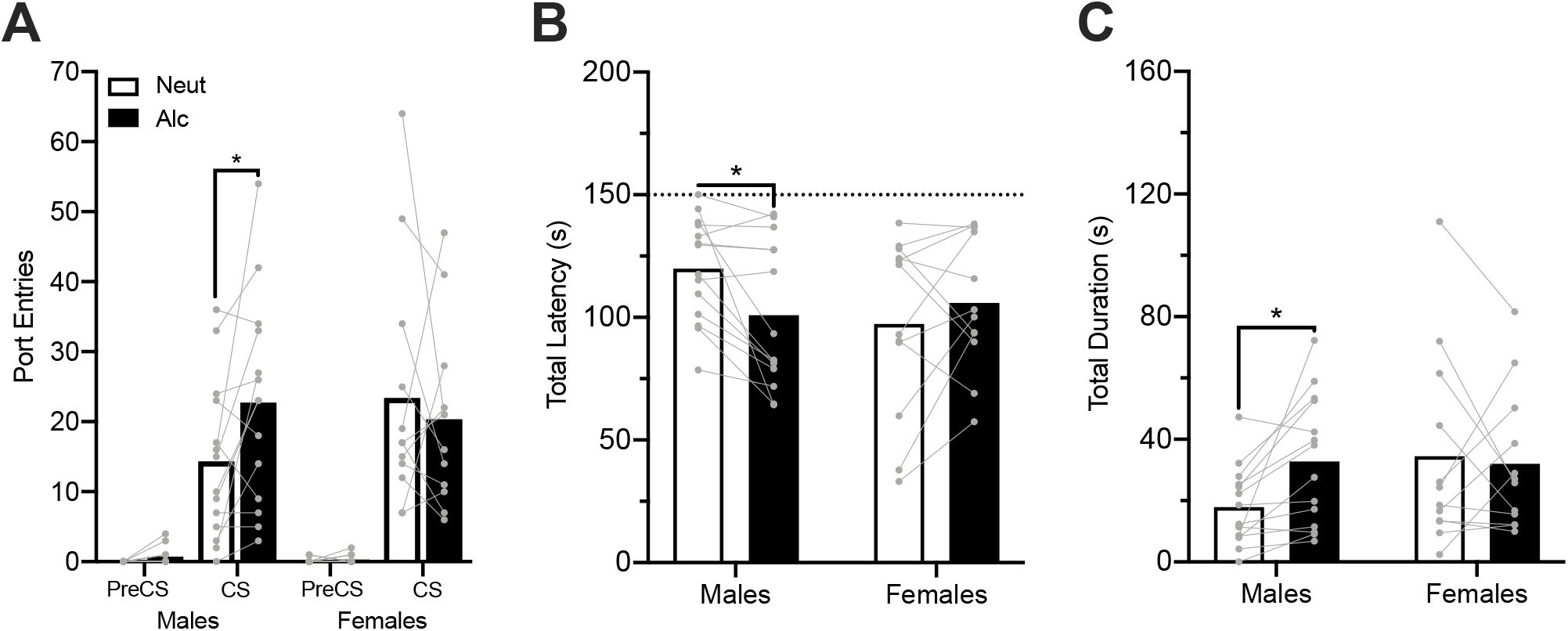
CS port entries at test, in the alcohol context (filled bars) and neutral context (open bars) in the absence of ethanol delivery, in males and females. Grey circles depict data from individual rats. \**p* < 0.05. (**A**) Mean PreCS and CS port entries. CS port entries were significantly greater in the alcohol context than in the neutral context, in males. (**B**) Mean latency to alcohol CS port entries. The latency to initiate CS port entries was significantly shorter in the neutral context compared to the neutral context, in males. (**C**) Mean total duration of port entries initiated during the CS. Total duration of CS port entries was significantly greater in the alcohol context than in the neutral context, in males.

Our prior research has reliably found that male rats make more CS port-entries at test in the alcohol context than in the neutral context (Millan et al., 2015; Remedios et al., 2014; Sciascia et al., 2015; Valyear et al., 2020). Visual inspection of Figure 3A and the trending Context x Sex interaction suggested that we replicated this result in males, but that females did not show a context-dependent modulation of CS port entries. Planned comparison of CS port-entries revealed that males made more CS port-entries in the alcohol context (*M* = 22.71 ± *SE* = 3.96) than in the neutral context (*M* = 14.29 ± *SE* = 3.03) [*t*_(13)_ =2.50, *p*=.027, *d*=.668, 95% CI (0.08, 1.24)], while CS port-entries in females did not differ between the alcohol context (*M* = 20.33 ± *SE* = 3.73) and the neutral context (*M* = 17.44 ± *SE* = 5.04) [*t*_(11)_ =-0.56, *p*=.584, *d*=-.16, 95% CI (−0.73, 0.41)].

The total latency to initiate port-entries after CS onset (Figure 4B) was shorter in the alcohol context compared to the neutral context in males but was similar across contexts in females. Overall latency did not vary across contexts [Context, *F*_(1,24)_=1.01, *p*=.326, *η*^*2*^=.008] or sex [Sex, *F*_(1,24)_=0.75, *p*=.396, *η*^*2*^=.030]. However, latency of port-entries initiated during the CS varied as a function of context across sex [Context x Sex, *F*_(1,24)_=6.97, *p*=.014, *η*^*2*^=.055]. Planned comparisons using paired samples t-tests revealed that males were faster to initiate CS port-entries in the alcohol context (*M* = 100.939 ± *SE* = 7.927) compared to the neutral context (*M* = 119.84 ± *SE* = 5.68) [*t*_(13)_=-3.00, *p*=.011, *d*=-.80, 95% CI (−1.39, −0.18)]. However, the time it took in females to initiate the CS port-entries did not differ between the alcohol context (*M* = 105.86 ± *SE* = 7.89) and the neutral context (*M* = 97.37 ± *SE* = 10.59) [*t*_(13)_=1.01, *p*=.335, *d*=.291, 95% CI (−0.29, 0.86)].

The total duration of the port-entries initiated during the CS (Figure 4C) was greater in the alcohol context compared to the neutral context in males, while in females the total duration of CS port-entries did not differ across contexts. Overall, the total duration of CS port-entries did not differ across context or sex [Context, *F*_(1,24)_=1.91, *p*=.180, *η*^*2*^=.018; Sex, *F*_(1,24)_=1.01, *p*=.325, *η*^*2*^=.040; Context x Sex, *F*_(1,24)_=3.73, *p*=.065, *η*^*2*^=.035]. Planned comparisons using paired samples t-tests found that in males the total duration of CS port-entries was greater in the alcohol context (*M* = 32.794, *SE* = 5.688) than in the neutral context (*M* = 17.94 ± *SE* = 3.43) [*t*_(13)_=3.11, *p*=.008, *d*=.831, 95% CI (0.21, 1.43)], whereas in females the total duration CS-elicited port-entries was similar in the alcohol context (*M* = 31.99 ± *SE* = 6.62) and the neutral context (*M* = 34.46 ± *SE* = 9.29) [*t*_(11)_=-0.31, *p*=.762, *d*=-.090, 95% CI (−0.66, 0.48)].

### Alcohol-Priming Induced Reinstatement Test

Following repeated tests sessions an alcohol-primed reinstatement test was conducted, in which 0.2 ml of ethanol was delivered at the start of the session and during the first CS presentation. Compared to the extinction baseline (Figure S1), CS-elicited port-entries were reinstated in both the alcohol context and neutral context, in males and females. However, CS port-entries at test were higher in the alcohol context compared to the neutral context in males but not females (Figure 6A). Overall, ΔCS port-entries were higher during the reinstatement test than the extinction baseline [Phase, *F*_(1,22)_=81.48, *p*<.001, *η*^*2*^=.527] and were greater in the alcohol context than in the neutral context [Context, *F*_(1,22)_=7.87, *p*=.010, *η*^*2*^=.073; Phase x Context, *F*_(1,22)_=4.70, *p*=.041, *η*^*2*^=.030]. No other differences were found [Sex, *F*_(1,22)_=1.99, *p*=.173, *η*^*2*^=.018; Phase x Sex, *F*_(1,22)_=0.52 *p*=.477, *η*^*2*^=.003; Context x Sex, *F*_(1,22)_=.02, *p*=.891, *η*^*2*^<.001; Phase x Context x Sex, *F*_(1,22)_=0.39, *p*=.538, *η*^*2*^=.003].

**Fig. 6.**
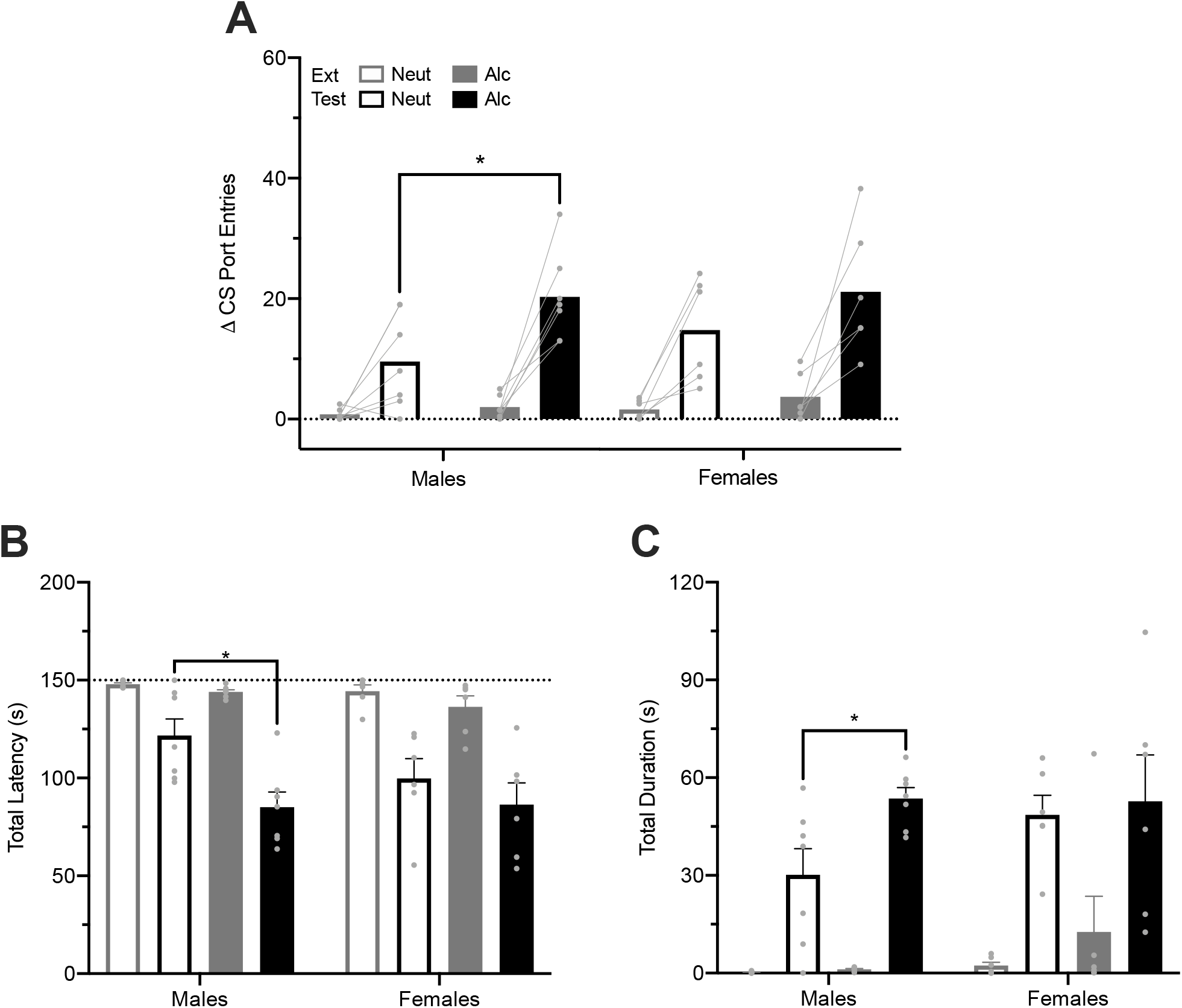
Port entries in the neutral context (empty bars) and alcohol context (filled bars) during alcohol-primed reinstatement test (black) compared to the extinction baseline (average of last two extinction sessions; grey), in males (left panels) and females (right panels). Circles depict data from individual rats. \**p* < 0.05. (**A**) Mean (± SEM) ΔCS port entries (CS minus PreCS por entries). ΔCS port entries at test were significantly greater in the alcohol context compared to the neutral context, in males. (**B**) Mean (± SEM) total latency to initiate CS port entries. The latency to initiate CS port entries was significantly shorter in the alcohol context than in the neutral context, in males. (**C**) Mean (± SEM) total duration of port entries initiated during the CS. The total duration of CS port entries was significantly greater in the alcohol context than in the neutral context, in males.

Our prior research found that male rats made more CS port-entries in the alcohol context than in the neutral context during reinstatement (Valyear et al., 2020). Visual inspection of Figure 6A suggested that despite the lack of a significant three-way Phase x Context x Sex interaction, this effect may have been replicated in males but not in females. Planned comparisons revealed that males made more CS port-entries at test in the alcohol context (M = 20.29 ± *SE* = 2.78) than in the neutral context (*M* = 9.57 ± *SE* = 2.95) [*t*_(12)_=-2.65, *p*=.021, *d*=-1.41, 95% CI (−2.58,-0.20)]. However, in females CS port-entries at test were similar in the alcohol (*M* = 21.00 ± *SE* = 4.26) and neutral (*M* = 14.67 ± *SE* = 3.49) contexts [*t*_(10)_=-1.13, *p*=.283, *d*=-0.66, 95% CI (−1.81, 0.53)].

The total latency to initiate port-entries following CS onset was shorter in the test than the extinction baseline and was shorter in the alcohol context compared to the neutral context in males, but not in females (Figure 6B). Overall total latency was shorter during the test compared to the extinction baseline [Phase, *F*_(1,22)_=105.11, *p*<.001, *η*^*2*^=.570], was shorter in the alcohol context compared to the neutral context [Context, *F*_(1,22)_=8.28, *p*=.009, *η*^*2*^=.067; Phase x Context, *F*_(1,22)_=4.69, *p*=.041, *η*^*2*^=.025]. No sex differences were found [Sex, *F*_(1,22)_=2.22, *p*=.151, *η*^*2*^=.018; Phase x Sex, *F*_(1,22)_=0.29, *p*=.593, *η*^*2*^=.002; Context x Sex, *F*_(1,22)_=0.79, *p*=.384, *η*^*2*^=.024; Context x Phase x Sex, *F*_(1,22)_=2.41, *p*=.135, *η*^*2*^=.013]. However, planned comparisons indicated that the total latency to initiate CS port-entries at test was shorter in the alcohol context (*M* = 85.15 ± *SE* = 7.66) than the neutral context (*M* = 121.71 ± *SE* = 8.53) in males [*t*_(12)_=3.19, *p*=.008, *d*=1.71, 95% CI (0.35, 2.80)]. In contrast, the total latency to initiate CS port-entries at test did not differ between the alcohol (*M* = 86.35 ± *SE* = 11.19) and the neutral (*M* = 99.74 ± *SE* = 10.17) contexts in females [*t*_(10)_=0.89, *p*=.397, *d*=0.511, 95% CI (−0.69,1.61)].

The total duration of port-entries initiated during the CS was greater during the test compared to the extinction baseline and was greater in the alcohol context than in the neutral context in males, but not females (Figure 6C). Overall, the total duration of CS port-entries at test was greater than at the extinction baseline [Phase, *F*_(1,22)_=93.30, *p*<.001, *η*^*2*^=.630]. No other effects were found [Context, *F*_(1,22)_=0.79, *p*=.383, *η*^*2*^=.007; Phase x Context, *F*_(1,22)_=0.21, *p*=.650, *η*^*2*^=.001; Sex, *F*_(1,22)_=0.91, *p*=.350, *η*^*2*^=.008; Phase x Sex, *F*_(1,22)_=0.06, *p*=.806, *η*^*2*^<.001; Context x Sex, *F*_(1,22)_=0.07, *p*=.790, *η*^*2*^<.001; Phase x Context x Sex, *F*_(1,22)_=0.77, *p*=.389, *η*^*2*^=.005]. Planned comparisons revealed that the total duration of CS port-entries at test were greater in the alcohol context (*M* = 53.59 ± *SE* = 3.34) relative to the neutral context (*M* = 30.21 ± *SE* = 8.01) in males [*t*_(8.03)_=-2.70, *p*=.020, *d*=-1.44, 95% CI (−2.55, −0.09)]. In females, the total duration of CS port-entries at test was similar in both the alcohol (*M* = 52.76 ± *SE* = 14.25) and neutral (*M* = 48.57 ± *SE* = 6.00) contexts [*t*_(10)_=-0.27, *p*=.792, *d*=-0.16, 95% CI (−1.29, 0.98)].

## DISCUSSION

The current study investigated potential sex differences in the influence of an alcohol-associated context on responding to an alcohol-predictive CS. We found that CS port-entries in males were invigorated in the alcohol context, compared to the neutral context. In contrast, CS port-entries at test in females were similar in both contexts. In males, alcohol-priming induced reinstatement of CS port-entries was higher in the alcohol context compared to the neutral context, while in females this effect was similar in both the alcohol and neutral contexts. Our findings support the notion that context enhances conditioned responding in males but not in females.

Males and females did not differ in ingested dose of ethanol or ethanol consumption in the home-cage ethanol exposure. Commonly, females have been shown to consume more ethanol per bodyweight compared to males in this procedure, although results vary depending on the strain of rat used, drinking conditions, and experimental procedures (Hilderbrand and Lasek, 2018; Priddy et al., 2017). The ingested dose of ethanol in males from the current study align with previous findings from our laboratory using the same rat strain, showing similar ingested dose of ethanol (Supplementary Material; Valyear et al., 2020). It is possible that the strain of rat contributed to the lower levels of ethanol consumption in both sexes. Furthermore, males weighed more than females, and although both sexes consumed similar amounts of ethanol, males also consumed more water than females. These results may suggest that females had higher blood alcohol concentrations (BAC) compared to males, which should be investigated in future work. In humans, women have been shown to reach higher peak BAC and are more susceptible to alcohol’s pharmacological effects compared to men after consuming the same doses (Mumenthaler et al., 1999).

In the Pavlovian discrimination training phase rats were trained to discriminate between an alcohol-associated context in which a CS was paired with alcohol delivery and a second neutral context in which a NS was presented without alcohol delivery. Both sexes were able to discriminate between the alcohol and neutral contexts as total port-entries remained high in the alcohol context and decreased in the neutral context throughout the sessions. Overall, females entered the fluid port more than males. There were no sex differences during non-CS time interval such as the ITI (Figure S1), suggesting that this effect was driven by the females making more port-entries during the CS. Both sexes displayed an increase in CS port-entries in the alcohol context compared to the low amount of NS port-entries in the neutral context across training sessions, with females making more CS port-entries in the alcohol context than males. It may be the case that females acquired the alcohol context or cue association more effectively than males. Previous findings of females acquiring and retaining associations more effectively than males have been shown in associative learning paradigms (Dalla and Shors, 2009). Females have also been shown to make more entries into a dipper containing alcohol compared to males during operant self-administration (Nieto and Kosten, 2017). Notably the total durations of CS port-entries and the latency to produce CS port-entries did not differ across sex, suggesting that both males and females were equally motivated to make port-entries during the CS.

At test, the CS was presented in both the alcohol and neutral contexts, without alcohol delivery. We found that in males, CS port-entries were significantly invigorated in the alcohol context compared to the neutral context. In addition, males made longer CS port-entries and were faster to initiate CS port-entries in the alcohol context compared to the neutral context. Thus, an alcohol-associated context modulates conditioned responding to a alcohol-predictive CS in males. This finding replicates previous findings from our laboratory using male rats (Sciascia et al., 2015; Valyear et al., 2020) and supports the notion that the context in which a drug is consumed can influence responding to a discrete drug predictive CS in males (Chaudhri et al., 2009; Conklin, 2006; Crombag and Shaham, 2002; Janak, 2013; Remedios et al., 2014; Zironi et al., 2006). In females, the number of CS port-entries, total latency, and total duration was similar in both contexts. Thus, females responded to a discrete alcohol-predictive CS in a context-independent manner at test. As both males and females were able to acquire the association between context and stimuli, the observed sex difference was not due to differential learning during training. This pattern of conditioned responding in females has also been shown in renewal procedures, which have revealed that conditioned responding of females in the experimental (i.e. ABA) and control (i.e. AAA) groups does not differ at test. Males in the experimental group respond more at test compared to the control group (Anderson and Petrovich, 2015, 2018a, 2018b). Therefore, context may modulate CS port-entries in males, but not in females.

A similar sex difference was found during the alcohol-primed reinstatement test, in which alcohol was presented at the start of the session and during the first CS presentation, in both contexts. In males, CS port-entries were reinstated in both contexts but were invigorated in the alcohol context compared to the neutral context. This finding replicates previous literature showing that reinstatement of responding to a discrete alcohol-predictive CS is invigorated in a context associated with alcohol, in male rats (Valyear et al., 2020). Males in the alcohol context were also faster to initiate CS port-entries and made longer CS port-entries in the alcohol context compared to the neutral context, suggesting that in the alcohol context rats were more motivated to produce port-entries during the CS. In females, CS port-entries were also reinstated in both contexts, but CS port-entries, latency, and duration did not differ across the contexts. Given that no sex differences were found during the repeated tests and responding in both sexes significantly decreased throughout the sessions in both contexts (Figure S2), this effect cannot be explained by the lack of extinction. There has been evidence to suggest that drug-associated contexts do not impact reinstatement of cue-elicited responding in females. For example, in context-driven reinstatement of methamphetamine self-administration, males responded more to the methamphetamine-paired lever than to the inactive lever, while females responded similarly to both levers, in the drug associated context (Takashima et al., 2018). Thus, context may modulate alcohol-priming induced reinstatement in males but not in females.

The sex differences found during the tests in the current study, are supported by the notion that males and females utilize contextual information differently. In studies using spatial tasks that manipulate both landmarks within the environment and the environmental context, men predominately rely on the contextual cues to solve the task. In contrast, women predominately rely on landmark cues to guide their behaviour in these tasks (Chai and Jacobs, 2010; Keeley et al., 2013; Sandstrom et al., 1998; Saucier et al., 2002; Shah et al., 2013). Similar findings in preclinical studies have shown that males and females rely on different types of environmental cues (e.g. distal or proximal) to complete spatial tasks (Rodríguez et al., 2011; Tropp and Markus, 2001; Williams et al., 1990; Williams and Meck, 1991). Therefore, males may be more reliant on environmental information, while females are more reliant on cues than the surrounding environment to guide behaviour. In the context of the current study, this may indicate that male rats may have relied on the context as a predictor of alcohol, while females may have relied on the CS as a predictor of alcohol.

The acquisition of contextual representations may also differ across sex. For example, females have lower levels of freezing in a context associated with shock compared to males, but this difference is not evident when both sexes are pre-exposed to the context (Wiltgen et al., 2001). Furthermore, in contextual fear conditioning, females are more likely to generalize fear to neutral contexts compared to males, but this effect is diminished following context preexposure (Asok et al., 2019; Keiser et al., 2017). Notably, females have been shown to generalize fear to similar contexts with training, suggesting a decrease of context specificity with time (Lynch et al., 2013). Thus, females may process contextual information differently, and may require more training using distinct contexts compared to males. However, whether these effects contributed to the sex differences found in the current study remains to be tested.

The differential learning and utilization of contextual information to predict a drug of abuse can have great implications on treatment in clinical settings. In humans, the gap in alcohol consumption between men and women is narrowing, especially with age (Keyes et al., 2019). Additionally, women with alcohol use-disorder are at greater risk of alcohol-related deaths, brain and organ damage, cognitive dysfunctions, and co-morbid mental health disorders (Ceylan-Isik et al., 2010; Walter et al., 2003). Therefore, it may be the case that drug craving and the risk of relapse is driven by different learning and neurobiological mechanisms in men and women. Due to the large discrepancy between men and women in the physiological, psychological, behavioural, and neurobiological factors involved in drug use (Becker et al., 2017), sex differences need to be taken into account in assessment and treatment of substance use disorders. The need for considerations based on sex, is further highlighted by studies using gender-specific treatment programs for drug addiction that have shown better outcomes than mixed-gender programs (Polak et al., 2015).

In conclusion, these findings suggest that the modulatory influence of an alcohol-associated context on an alcohol-predictive CS operates as a function of biological sex. We replicated findings in males, showing that they made more CS port-entries in the alcohol-associated context than in the neutral context at test and during an alcohol-primed reinstatement test. Females made similar number of CS port-entries in both the alcohol-associated context and the neutral context at both tests, suggesting that their responding was not modulated by context. These data identify novel sex differences in the capacity of a drug-associated context to influence cue-elicited responding and may indicate potential sex differences in context utilization and processing.

## ACKNOWLEDGMENTS

This research was funded by the Canadian Institution of Health Research (MOP-137030, N.C.). N.C. is a member of the Center for Studies in Behavioral Neurobiology. D.S. is supported by a graduate bursary from the Center for Studies in Behavioural Neurobiology at Concordia University. The authors would like to thank Mohammed Akhtar for his technical assistance, João Vitor de Camargo for assistance with reviewing literature, and Stephen Cabilio for support with Med-PC programming.

## Notes

### Competing Interest Statement

The authors have declared no competing interest.

