## Supplementary Material for "Sex differences in the amplification of responding to an alcohol-predictive cue by an alcohol-associated context"

Rat Breeding Information

The rats used in the current study were bred in our on-site TH::Cre rat breeding colony, in which 4 heterozygote TH::Cre positive dams were paired with 2 wild type sires (Charles River, St. Hubert, QC). Pups were weaned at twenty-three days post-natal, seprated by sex, and pair-housed in polycarbonate shoebox cages containing beta chip bedding (Aspen Sani chips; Envigo, Indianapolis, IN), a nylabone chew toy (Nylabones, Bio-Serv, Flemington, NJ), a red plastic tunnel (Rat retreats, Bio-Serv, Flemington, NJ), and shredded paper for enrichment, with unrestricted access to standard rat chow (Rodent Diet, Charles River, St. Hubert, QC) and water. Males and females were housed on separate walls within the breeding colony room held at 21 +/- 2°C and approximately 40-50% humidity on a 12-hour light-dark cycle (lights on at 0700 hours). One week following weaning, two rounds of pinna samples were collected, and two rounds of genotyping were conducted to verify the genotype of each rat. Those that showed a negative genotype in both rounds (n = 31) were selected for the current study. Rats were transferred into the colony room at ~9 weeks of age and single housed (See Methods).

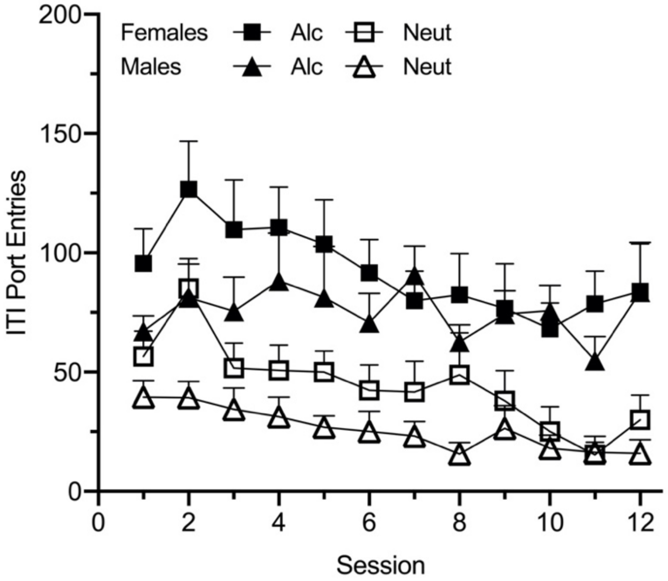


**Figure S1.** Port entries in the Pavlovian discrimination training initiated during the variable inter-trial interval (ITI), in males (triangles) and females (squares) in the alcohol (filled shapes) and neutral (empty shapes) contexts. ITI port entries varied as a function of session [Session, $F(5.098, 122.346) = 4.257$, $p = .001$, $\eta^2 = .046$] and was greater in the alcohol context than the neutral context [Context, $F(1,24) = 67.462$, $p < .001$, $\eta^2 = .205$], no other interactions or sex differences [Context x Session; $F(6.338, 152.100) = 0.891$, $p = .508$, $\eta^2 = .006$, Context x Sex; $F(1,24) = 0.06$, $p = .808$, $\eta^2 < .001$, Session x Sex; $F(5.098, 122.346) = 1.451$, $p = .210$, $\eta^2 = .016$, Context x Session x Sex; $F(6.338, 152.100) = .735$, $p = .629$, $\eta^2 = .005$, Sex; $F(1,24) = 3.525$, $p = .073$, $\eta^2 = .029$].

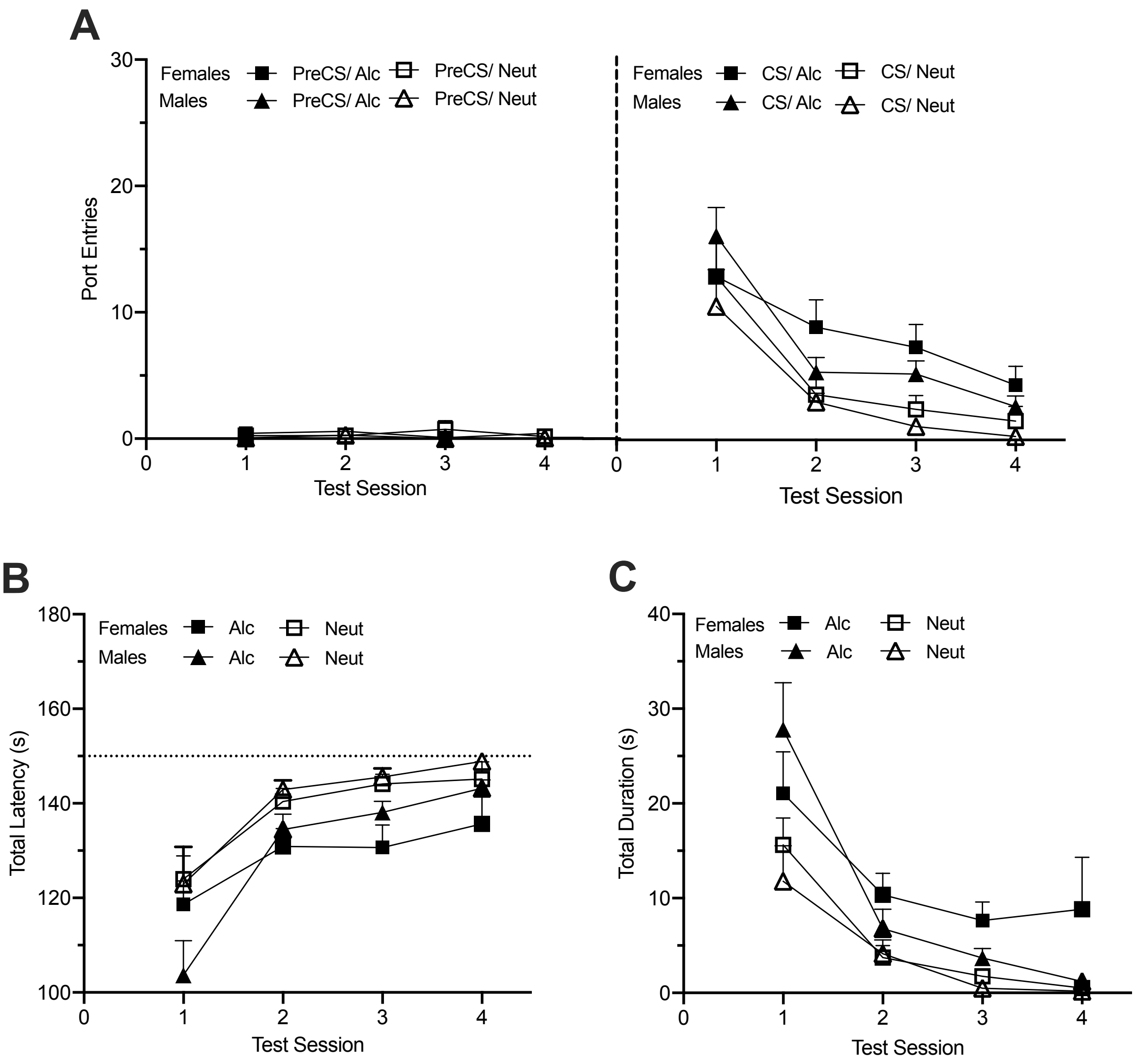


**Figure S2.** Extinction of CS port entries in the alcohol context (filled shapes) and neutral context (empty shapes) in the absence of alcohol delivery during repeated tests, in males (triangles) and females (squares).

(**A**) Mean (± SEM) port entries during the PreCS (left panel) and CS (right panel) intervals in the alcohol and neutral contexts. Port entries varied as a function of session [Session, $F(1.98, 47.4) = 41.13$, $p < .001$, $\eta^2 = .120$], were higher in the alcohol context than in the neutral context [Context, $F(1,24) = 16.94$, $p < .001$, $\eta^2 = .020$], and varied as a function of interval [Interval, $F(1,24) = 117.31$, $p < .001$, $\eta^2 = .239$; Interval x Context, $F(1,24) = 17.30$, $p < .001$, $\eta^2 = .020$]. Across sessions, CS port entries decreased, whereas PreCS port entries remained low [Interval x Session, $F(1.89, 45.35) = 47.95$, $p < .001$, $\eta^2 = .120$] with no difference as a function of context [Context x Session, $F(1.74, 41.73) = 0.22$, $p = .777$ $\eta^2 < .001$]. The three-way Interval x Context x Session interaction was not significant [$F(1.92, 46.11) = 0.41$, $p = .776$, $\eta^2 = .001$]. No sex differences were found [Sex, $F(1,24) = 1.59$, $p = .220$, $\eta^2 = .004$; Interval x Sex, $F(1,24) = .71$, $p = .408$, $\eta^2 = .001$; Session x Sex, $F(1.98, 47.39) = 0.65$, $p = .525$, $\eta^2 = .002$; Context x Sex, $F(1,24) = 0.03$, $p = .876$, $\eta^2 < .001$; Interval x Session x Sex, $F(1.89, 45.35) = 0.63$, $p =$

.531, $\eta^2$ = .002; Interval x Context x Sex, $F(1,24) = 0.06$, $p = .807$, $\eta^2 < .001$; Interval x Context x Session x Sex, $F(1.92, 46.11) = 1.30$, $p = .283$, $\eta^2 = .005$].

(**B**) Mean (± SEM) total latency (s) to initiate port entries during the CS in the alcohol and neutral contexts. The total latency to initiate a CS port entry varied as a function of session [Session, $F(1.57, 37.57) = 36.44$, $p < .001$, $\eta^2 = 284$] and was greater in the alcohol context than in the neutral context [Context, $F(1,24) = 25.64$, $p < .001$, $\eta^2 = .068$] with no difference across sessions [Context x Session, $F(2.06, 49.47) = 0.26$, $p = .779$, $\eta^2 = .002$]. No sex differences were found [Sex, $F(1,24) = 0.17$, $p = .682$, $\eta^2 = .007$; Session x Sex, $F(1.57, 37.57) = 2.67$, $p = .094$, $\eta^2 = .021$; Context x Session x Sex, $F(2.06, 49.47) = 1.27$, $p = .291$, $\eta^2 = .011$].

(**C**) Mean (± SEM) total duration of port entries initiated during the CS in the alcohol and neutral contexts. The total duration of CS port entries varied across sessions [Session, $F(1.63, 39.23) = 34.42$, $p < .001$, $\eta^2 = .304$] and was greater in the alcohol context than in the neutral context [Context, $F(1,24) = 21.21$, $p < .001$, $\eta^2 = .066$]. The two-way Context x Session interaction was not significant [$F(1.99, 47.83) = 1.40$, $p = .256$, $\eta^2 = .012$]. No sex differences were found [Sex, $F(1,24) = 1.29$, $p = .267$, $\eta^2 = .005$; Session x Sex, $F(1.63, 39.23) = 0.79$, $p = .438$, $\eta^2 = .007$; Context x Sex, $F(1,24) = 0.10$, $p = .757$, $\eta^2 < .001$; Context x Session x Sex, $F(1.99, 47.83) = 2.32$, $p = .110$, $\eta^2 = .020$].
